# Effect of acute elevated magnesium on bursting activity and information-processing dynamics in cortical cultures

**DOI:** 10.1101/2025.08.20.671363

**Authors:** Kathryn G. H. LaGrange, Thomas F. Varley, Lilly F. O’Shea, Julia G. Fitzpatrick, Sarah E. Cummings, John M. Beggs

## Abstract

Magnesium (Mg^2+^) plays a significant role in hippocampal memory and learning and is implicated in a variety of neurological disorders, such as migraine. Despite this crucial role Mg^2+^ has on brain health, its effect on the dynamics of networks of neurons is still not fully understood. This study investigates the impact of several doses of elevated extracellular Mg^2+^ on cortical **organotypic cultures**. Cultures were recorded on a 512-microelectrode array and analyzed using the burstiness index (BI), the rate of network-wide bursts relative to other neuronal activity. We also use a suite of information theoretic measures to further establish the role Mg^2+^ plays in the brain. Elevated Mg^2+^ is found to have a dose-dependent increase on BI caused by a loss of network activity not contained within a burst or high firing rate activity. Information dynamics further show that the network experiences a loss of entropy and an increase in time-reversibility, connectivity, and transfer of redundant information. These factors indicate that Mg^2+^ causes a loss of complex activity and an increase in highly integrated, constant dynamics. This lays the groundwork for a deeper examination of the role Mg^2+^ plays in learning, memory and treating neurological disorders.

**AUTHOR SUMMARY:** Magnesium is crucial for neurological health. In pyramidal neurons, magnesium blocks NMDA receptors, thereby reducing synchronous neural activity in the brain. This has led to the canonical view of magnesium as a “burst quieter”. Here, we take a closer look at magnesium’s role in cortical slice cultures by examining network burstiness with normal and chronically elevated levels of magnesium. We show that magnesium can effectively increase burstiness, not reduce it. Magnesium is further shown to cause significant changes in how neurons process information. These findings may have impact how magnesium is used as a treatment for neurological disorders.

## INTRODUCTION

Magnesium (Mg^2+^) is a metallic cation that plays a key role in health throughout the body, including in the brain de Baaij, Hoenderop, and Bindels (2015). Maintaining a healthy balance of Mg^2+^ is important for cognitive performance and neurological health. Elevated dietary Mg^2+^ has been shown to improve performance in hippocampal tasks involving learning and spatial working, short- and long-term memory Slutsky et al. (2010). A dietary Mg^2+^ deficiency decreases performance in such hippocampal tasks, including spatial memory, social memory and contextual fear Serita et al. (2019). Low levels of Mg^2+^ have also been associated with neurological disorders including migraine Dolati, Rikhtegar, Mehdizadeh, and Yousefi (2020) and Alzheimer’s disease Chui et al. (2011) and may be related to chronic pain, epilepsy, Parkinson’s disease, anxiety and depression Kirkland, Sarlo, and Holton (2018). Despite elevated Mg^2+^ being a proposed treatment for some of these disorders (e.g. Dolati et al. (2020)), it is still not fully understood what effect it has on neuronal network functions.

On the receptor scale, Mg^2+^ plays a significant role in the activation of N-methyl-D-aspartate (NMDA) receptors. **NMDA receptors (NMDAR)** appear on approximately 80% of neurons in the cerebral cortex, but are most often associated with activation of excitatory pyramidal neurons Conti (1997). NMDAR are voltage-gated Ca^2+^ channels that act as coincidence detectors, requiring both the binding of glutamate and removal of a Mg^2+^ molecule to activate. This Mg^2+^ molecule is removed by a sufficiently large depolarization Mayer, Westbrook, and Guthrie (1984); Nowak, Bregestovski, Ascher, Herbet, and Prochiantz (1984). Increasing extracellular Mg^2+^ increases the firing threshold of pyramidal neurons, effectively disabling excitatory neurons.

One network-wide effect elevated Mg^2+^ has is a disruption of bursts. Bursts are network spiking events where a large number of neurons rapidly fire in a short period of time. At large time scales, these bursts may appear synchronous, but short time scales reveal a high level of complexity (for review, see Corner, van Pelt, Wolters, Baker, and Nuytinck (2002)). Bursts can last up to hundreds of milliseconds, but even those lasting only a couple milliseconds can be broken into repeating chains of activity with complex neuron-to-neuron interactions Beggs and Plenz (2003). Mg^2+^ acts as a “burst quieter” by disrupting this chain and preventing the ensuing network-wide activation. Bursts occur less frequently, thereby enhancing synaptic recovery Tabak, Mascagni, and Bertram (2010). This results in shorter, highly synchronized bursts Teppola, Aćimović, and Linne (2019) that are mediated by -amino-3-hydroxy-5-methyl-4-isoxazolepropionic acid (AMPA) receptors rather than NMDAR.

In its role as a burst quieter, elevated Mg^2+^ can increase network plasticity in neuronal cultures. Long term potentiation (LTP) is typically dependent on calcium (Ca^2+^) influx mediated by NMDAR Malenka and Nicoll (1993). Elevated Mg^2+^ has been applied in studies using electrical stimulation to trigger an artificial burst that results in network learning. Spontaneous network bursting can override the effects of this stimulation and reduce plasticity Maeda, Robinson, and Kawana (1995); Robinson et al. (1993); Wagenaar, Pine, and Potter (2006b). In such experiments, elevated Mg^2+^ quiets spontaneous bursts.

Stimulation provides a sufficiently high depolarization to surpass the Mg^2+^ block, allowing only stimulation-induced bursts to activate and affect network plasticity Maeda, Kuroda, Robinson, and Kawana (1998). A very high chronic dose of 12.5 mM Mg^2+^ has been shown to stop all spontaneous activity despite enhanced synaptic recovery and exclusively allow spikes caused by stimulation Baker, Ruijter, and Bingmann (1991), though with significant rates of cell death. At lower doses, the amount of Mg^2+^ required to stop bursting varies with the age of the culture Maeda et al. (1995).

In addition to quieting bursts, Mg^2+^ concentration impacts information flow within the cortex. NMDAR are known to have a significant effect on information flow through cortical layers. In adult animals, most NMDAR are found in the upper layers II/III of the cortex Currie, Wang, and Daw (1994). Projections from layer II/III to layer V are largely mediated by NMDAR Jones and Baughman (1988). Mg^2+^ concentration has been shown to affect this pathway Currie et al. (1994). It is likely that information transfer between small populations of cortical neurons is also affected by Mg^2+^ concentration, but this has not been previously investigated to our knowledge.

One of the core questions in neuroscience is how changes at the level of individual neurons (activation of receptors, changes in membrane polarization, etc.) produce global changes in patterns of activity across the whole system. One increasingly popular framework for studying the structure of these part-whole relationships is that of information dynamics Lizier (2013, 2014). Information dynamics uses information theory to decompose the structure of multipartite interactions into distinct “modes”: information storage, information transfer, and information integration/modification, which are roughly analogous to the major operations performed by a digital computer. This approach has become particularly popular in the study of spiking neural dynamics Newman, Varley, Parakkattu, Sherrill, and Beggs (2022), where discrete, binary spiking data is naturally suited to information-theoretic analyses.

Prior work has shown that information dynamics are sensitive to both behavioral Varley, Sporns, Schaffelhofer, Scherberger, and Dann (2023) and pharmacological Varley et al. (2024) differences, and here we adapt those approaches to study the effects of elevated Mg^2+^. Here we will provide a high-level, intuitive review of the various measures described here. For formal details, see the Materials and Methods section.

The most basic measure of the information content of a spiking neuron is the Shannon **entropy** of its spike train. Using a naive, maximum-likelihood estimator, the entropy is directly proportional to the spike rate. The entropy assumes each frame is a random sample from some binary probability distribution and has no temporal information. To assess how a single neuron “stores” information, we can use the **active information storage (AIS)** Wibral, Lizier, Vögler, Priesemann, and Galuske (2014). The AIS quantifies how much uncertainty about the *future* of a neuron is reduced by learning the state of its *past*. It can be thought of as an information-theoretic version of the autocorrelation function. The final first-order (i.e. single-neuron) measure is the **entropy production** Roldán and Parrondo (2010, 2012), which quantifies the difference between the forward and time-reversed trajectories. In systems that are far from equilibrium, spontaneous dynamics are irreversible. This leads to a production of entropy, which can be quantified by the Kullback-Leibler divergence between the time-forward and time-reversed state-transition structure. The entropy production, then, serves as a useful measure of “distance to equilibrium” for a given system.

To explore the structure of cell-cell interactions, we used the multivariate **transfer entropy** (mTE) Novelli and Lizier (2021); Novelli, Wollstadt, Mediano, Wibral, and Lizier (2019) to extract flows of information from a source neuron to a target neuron Schreiber (2000). The transfer entropy forms the second major component of the information dynamics framework after the active information storage. The distribution of information flows forms a directed, effective connectivity network, which can be characterized using techniques from network science Sporns (2010). The final measure of information dynamics is the information modification Lizier, Flecker, and Williams (2013), which quantifies the amount of “novel” information generated when a neuron performs a computation on multiple incoming streams Varley (2025). Unlike the active information storage and information transfer, information modification does not have a canonical form; here we follow Lizier (2013); Newman et al. (2022); Varley et al. (2024) and operationalize information modification with the synergistic information, as revealed by the /textbfpartial information decomposition (PID) Williams and Beer (2010). As part of the **synergy** computation, another higher-order information dynamic, the **redundancy**, is also provided “for free”.

Where the synergy represents the irreducible, integrated information produced by a target neuron, the redundancy is the information duplicated over all the source neurons.

Collectively, these three features (information storage, information transfer, and information modification) provide a quantitative scaffold on which to build a rigorous, mathematically precise understanding of how “computation” occurs in distributed, cortical circuits.

This study focuses on the impact an acute elevation of Mg^2+^ has on the network’s burst behavior and information dynamics. Organotypic cortical cultures were recorded on a microelectrode array (MEA) with control culture medium, multiple doses of elevated Mg^2+^ and then a washout of control medium. Contrary to the behavior in dissociated cultures, we found that organotypic cultures showed a significant dose-dependent increase in burstiness relative to other types of activity in the drug condition. To examine what may cause this difference, information dynamics between neurons were assessed in each recording condition.

## METHODS

### Organotypic Cultures

Organotypic cultures were prepared using the methods described in Tang et al. (2008) and Ito et al. (2011). All procedures followed guidelines from the National Institutes of Health and were approved by the Indiana University Animal Care and Use Committee, and all proper protocols for animal care were followed.

Briefly, neonatal P2-P6 Sprague-Dawley rats were sacrificed and their brains were extracted. The brains were coronally sliced using a vibratome to a thickness of 400 µm. The slices were placed on a permeable membrane with culture medium beneath them. They were incubated for two to four weeks at 37° C in a 95% O_2_, 5% CO_2_ concentration. Culture medium consisted of 1 L minimum essential medium (Sigma-Aldrich), 500 mL Hank’s balanced salt solution (Sigma-Aldrich), 500 mL heat inactivated horse serum (Sigma-Aldrich), 2 mL Penicillin-Streptomycin-Neomycin antibiotic mixture (Sigma-Aldrich) and 10 mL L-Glutamine (Sigma-Aldrich). The day after the dissection and then every three days, half of the culture medium was replaced to refeed the cultures.

### Recording

Following two to four weeks of incubation, the cortex of nineteen cultures was recorded on a 512-microelectrode array (MEA). The MEA has a triangular lattice of 5 µm diameter electrodes with 60 µm separating each electrode Litke et al. (2004). Data is recorded at a 20 kHz temporal resolution.

Throughout recording, culture medium is perfused at a rate of approximately 3 mL/minute. This culture medium is oxygenated with a mixture of 95% oxygen, 5% CO_2_ and kept heated to approximately the rat body temperature of 37° C.

Recording consisted of three one-hour conditions for each culture. During the first hour, control data was collected. A second batch of culture medium was then perfused that had an elevated MgCl_2_ concentration of 0 mM (sham), 1 mM, 3 mM or 5 mM. Recording resumed for this drug condition once the elevated Mg^2+^ medium was fully perfused. Finally, the control culture medium was fully perfused back to the slice and an hour-long washout condition was recorded. Spike sorting was completed with Kilosort 2.5 Pachitariu, Steinmetz, Kadir, Carandini, and D (2016) using a low-pass filter of 300 Hz and a custom map of the MEA.

### Burstiness Index

The **burstiness index (BI)** is an index representing a neuronal network’s tendency for all neurons to fire synchronously or independently Wagenaar, Madhavan, Pine, and Potter (2005). BI is a dimensionless index between 0 and 1, where 0 represents an even distribution of spike times and 1 represents an extreme tendency for all neuronal spikes to occur at the same times. To calculate the burstiness, each dataset was divided into 1 s time bins, as in Wagenaar et al. (2005). This long duration is enough to encapsulate an entire burst, which can have a duration of hundreds of milliseconds Baker, Corner, and van Pelt (2006). The total spikes were counted in each time bin, and the 15% of bins with the highest counts were determined. The choice of 15% was made following the observation that bursting will never occur in more than 15% of 1 s time bins within a given five minute recording Wagenaar et al. (2005). The percentage of all network spikes occurring within these bins *f*_15_ was used to calculate BI:

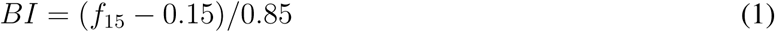

BI was calculated for each five minute window of recording, the same window as Wagenaar et al. (2005).

When interpreting BI, it is important to be mindful that it represents the proportion of spikes contained within network bursts to spikes outside of them. This results in two possible explanations for an increase in BI. First, BI increases any time more spikes are contained within bursts. This could be caused by an increase in burst frequency, burst duration or recruitment of more neurons during bursts. A second and less obvious possibility is that the burst behavior remains the same, but there is a reduction in the number of spikes outside of bursts. This would likewise increase the proportion of total spikes within bursts, leading to an increase in BI. Therefore, additional measures are required to interpret a change in BI.

### Dissociated Culture Data

Dissociated hippocampal culture data Timme et al. (2016) and dissociated neocortical culture data Wagenaar et al. (2005) were used from a previous study for comparison of BI values. Since 19 organotypic neocortical culture data sets were recorded in this study, 19 data sets were randomly selected from Timme et al. (2016) for comparison. Data was rebinned to 1 s for the calculation of BI. Since the full set of data for Wagenaar et al. (2005) was not available, surrogate data was created to complete statistical analysis. 19 surrogate BI values were randomly generated using a normal distribution. The surrogate values had the same mean and SEM as Wagenaar’s data (0.48 ± 0.02). To ensure the statistics were robust, thirty sets were generated and showed the same statistical agreement each time. Comparisons between organotypic and both types of dissociated culture BI were made using a Mann-Whitney U-test.

### Information Dynamics

In this paper, we replicate the methods described in Varley et al. (2024). We will provide the basic theory and parameters here and encourage interested readers to see the aforementioned literature.

*Shannon Entropy* The Shannon entropy was computed with a naive entropy estimator. Given the definition:

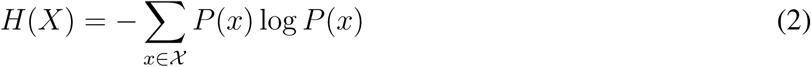

the probability *P* (*x*) was the number of times we observed *X* = *x* divided by the total number of samples. In this case, *X* could be on (spiking) or off (quiescent).

*Active Information Storage* For a temporal variable *X_t_*, the active information storage is quantifies as:

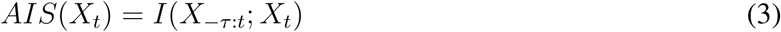

Where *τ* is the maximum history depth and *X*_−_*_τ_*:*_t_* is the joint-state of *X_t_*_−1_*, X_t_*_−2_*…X_t_*_−_*_τ_* . The AIS, then is the information that the past discloses about the next step *X_t_*. We used the IDTxl toolkit Wollstadt et al. (2019), which included an auto-embedding algorithm to determine the optimal embedding up to a maximum possible lag of 5. To control for variable entropy rates, we report the normalized *AIS*(*X_t_*)*/H*(*X_t_*).

*Entropy Production* The entropy production quantifies the degree of irreversibility in a neuron’s spike train. We can represent the forward and time-reversed version of *X_t_* as 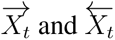 respectively. The entropy production is then diven by the Kullback-Leibler divergence between these two time series Roldán and Parrondo (2010, 2012):

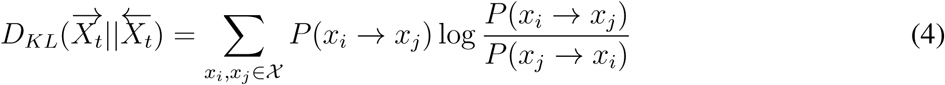

The larger this value, the more the time-forward statistics deviate from the time-reversed statistics and the farther the system is from equilibrium. Following Varley et al. (2024), we coarse-grained the 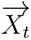 and 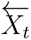 time series into 5 ms bins, to produce time series with 32 possible states, rather than a binary state.

This was done to ensure that the state-space was large enough to capture rich temporal dynamics. To satisfy the requirements of the Kullback-Leibler divergence, we only included forward and reverse transitions where both *P* (*x_i_* → *x_j_*) *>* 0 and *P* (*x_j_* → *x_i_*) *>* 0.

*Multivariate Transfer Entropy* The Shannon entropy, the active information storage, and the entropy production are all “low-order” measures: they describe the statistics of a single neuron. To understand how information flows from neuron to neuron, we use a multivariate extension of the classic transfer entropy Novelli and Lizier (2021); Novelli et al. (2019); Schreiber (2000). For a set of neurons **X** that collectively disclose information about a target *Y*, the multivariate transfer entropy is:

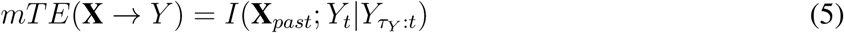

Here **X***_past_*refers to the joint, potentially multidimensional, embedding of every element in **X**, and *Y_τY_* _:*t*_ is the potentially multidimensional embedding of *Y* ’s past up to time *t* − 1.

The major benefit of the multivariate transfer entropy is that it accounts for synergies between multiple *X_i_*onto *Y*, as well as not double-counting redundancies (which can artificially inflate the apparent information-density of a system) Bossomaier, Barnett, Harré, and Lizier (2016). The catch, however, is that it is “higher-order”, in the sense that *mTE*(**X** → *Y*) is understood as something like a directed hyper-edge of multiple elements onto *Y* . To recover a pairwise network structure, links between a source *X^i^* and target *X^j^* are defined with the conditional mutual information: 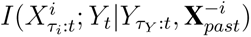.

For large systems with hundreds of neurons, it is impossible to fully optimize all possible source sets **X**, as well as optimal embeddings for every *X_i_* ∈ **X** and *Y* . Following Varley et al. (2024), we took a two-step filtering approach. First, for each target *Y*, we removed any prospective sources that had no significant bivariate transfer entropy over any lags on the range [-30,-1]. For efficient significance testing, we used the analytic null estimator based on the Chi-squared distribution Barnett and Bossomaier (2012), as implemented by the JIDT toolbox Lizier (2014). This restricted prospective parent set was then used to initialize the multivariate transfer entropy inference algorithm provided by the IDTxl toolbox Novelli and Lizier (2021); Wollstadt et al. (2019).

For consistency with prior work Ito et al. (2011); Nigam et al. (2016); Shimono and Beggs (2015); Varley et al. (2024), we further constrained the mTE inference algorithm to only consider one possible lag for each *X_i_* ∈ **X**: the lag that maximized the apparent transfer entropy. The optimal parent set **X** for each target neuron *Y* was inferred in parallel, and significance testing at every step of the inference pipeline was done using a set of 250 circular-shift nulls (designed to preserve first-order autocorrelations). Finally, the control for the effects of firing rate, we report normalized 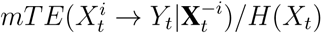.

*Network Measures* The multivariate transfer entropy inference produces an effective connectivity network, which describes how information flows from a source neuron *X^i^*to a target neuron *Y* in the context of all other parents of *Y* . To understand how Mg^2+^ modulated the global patterns of information flow, we characterized the topology of this network using standard statistics from network science. We considered the average *in-**degree*** (the average number of in-coming edges incident on a neuron), the average *out-degree* (the average number of out-going edges emerging from a neuron), and the **clustering coefficient** Holland and Leinhardt (1971); Watts and Strogatz (1998). The clustering coefficient for a given neuron quantifies what proportion of that neurons immediate neighbors are also neighbors themselves: a higher value indicates more closed triangles and greater “integration.” All network analyses were computed using the Igraph package in Python Csardi and Nepusz (2006).

*Redundancy and Synergy* The final measure of information dynamics is the information modification (also sometimes called “integration”). Here, following Lizier et al. (2013); Newman et al. (2022), we operationalize information modification using the synergistic information, as computed by the partial information decomposition (PID) Williams and Beer (2010).

The PID provides a decomposition of the joint mutual information 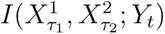 into four, non-overlapping “atomic” components (referred to as partial-information atoms):

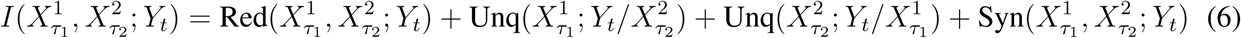

The redundant information, 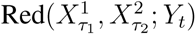 is the information about *Y_t_* that could be learned by observing 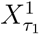 alone or 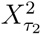 alone. The unique information 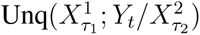 is the information about *Y_t_* that can only be learned by observing 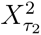 (in the context of 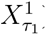). Finally, the synergistic information 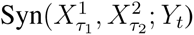 is the information about *Y_t_* that can only be learned when both sources are observed together. The synergistic information has been associated with “computation” and is a signature of causal colliders in multipartite systems Rosas, Mediano, and Gastpar (2024); Varley (2024).

The PID further requires the constraint that:

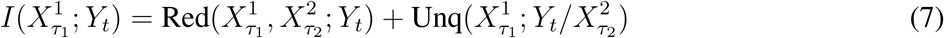

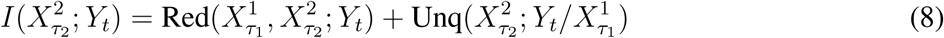

This produces a set of three known variables (the three mutual information) and four unknowns (the partial information atoms. If it is possible to specify the form of any atom, the remaining three are fixed. We used the *I_BROJA_* estimator of unique information Bertschinger, Rauh, Olbrich, Jost, and Ay (2014), as implemented in the IDTxl package Wollstadt et al. (2019). We computed the PID for all sets of motifs with two source neurons synapsing onto a single target, and fixed the lags *τ*_1_ and *τ*_2_ to be the same lags used in the multivariate transfer entropy network inference. Once again, all reported values are normalized by *H*(*X_t_*).

## RESULTS

### Burstiness

*Burstiness Index* The burstiness index (BI) was calculated for each network every five minutes. The average BI for all cultures did not show a trend over time (data not shown), so each five minute interval was included as a separate data point, as in Wagenaar et al. (2005). Figure 2 shows BI for the elevated Mg^2+^ (Figure 2*A*) and washout conditions (Figure 2*B*), where the groups are denoted by their concentration in the elevated Mg^2+^ condition. BI was normalized to the average BI baseline of the culture during the control condition. A linear regression showed no significant effect of concentration group in the control data (data not plotted, *R*^2^ = 6 ∗ 10^−5^*, p* = 0.90622). In the elevated Mg^2+^ condition, there was a significant positive correlation between normalized BI and dose (*R*^2^ = 0.025*, p <* 0.05). Washout with control medium removed this effect, showing no significant trend with dosage (*R*^2^ = 0.007*, p* = 0.283). The sham medium changes (0 mM Mg^2+^) yielded average normalized values at baseline in both Figure 2*A* and 2*B*, indicating that cultures did not experience an average change in BI in the absence of elevated Mg^2+^.

**Figure 1.**
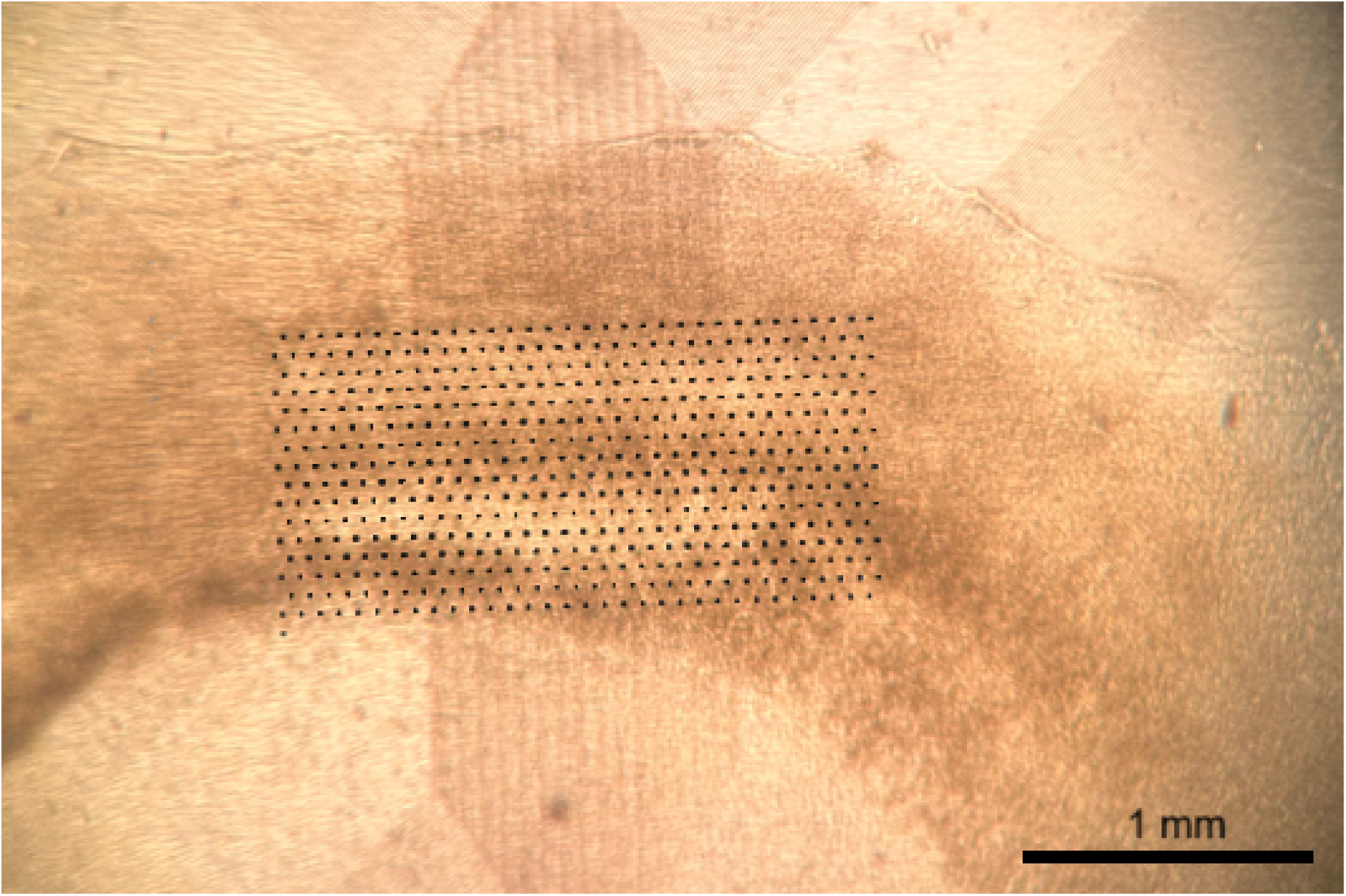
Sample organotypic culture on the electrode array. Cortical layers of the tissue are visible curving horizontally across the array. Slices were placed to record cortex. Black dots are superimposed on the image to show the positions of the electrodes.

**Figure 2.**
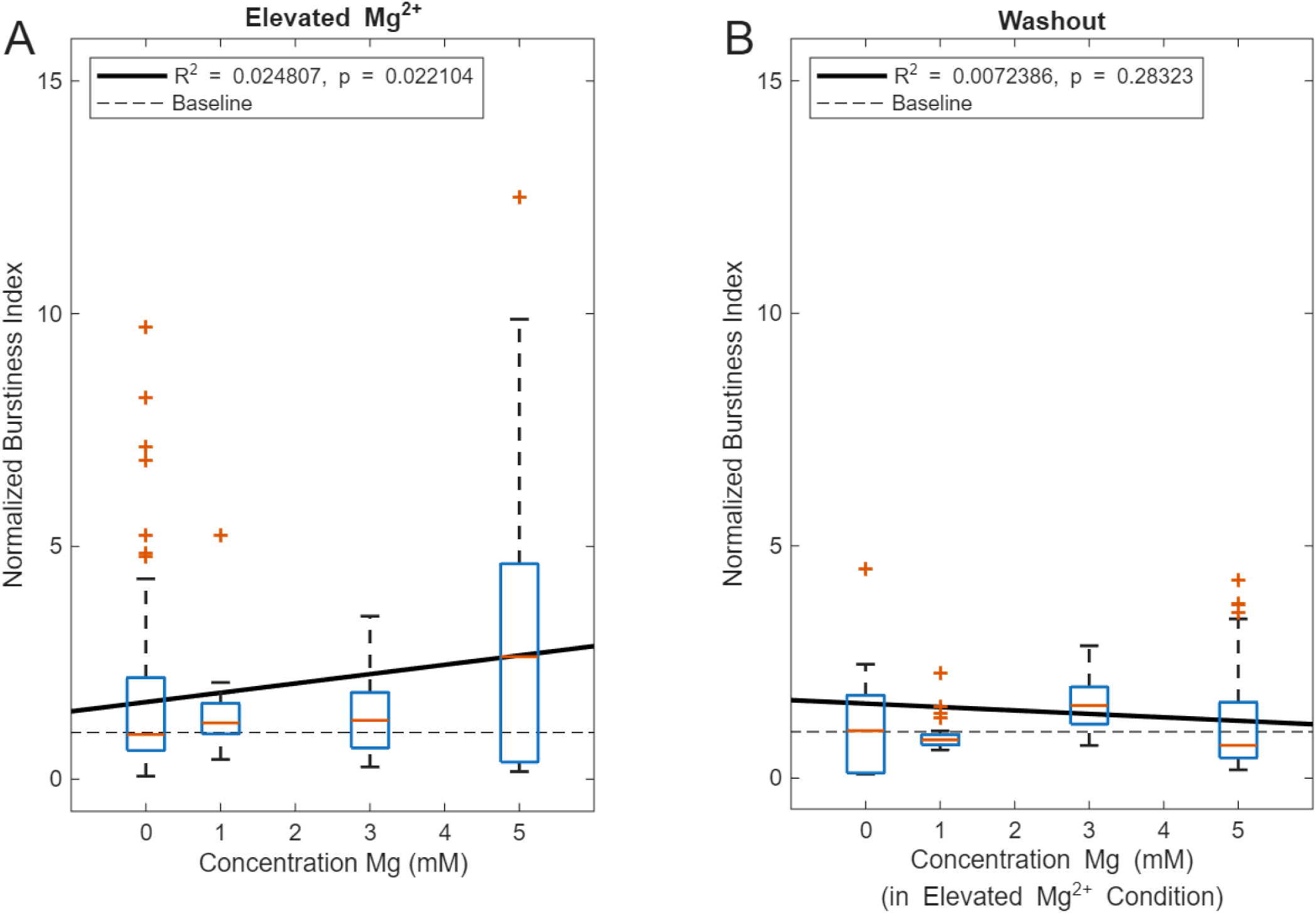
Elevated Mg^2+^ has a significant dose-dependent effect on the normalized Burstiness Index (BI). Normalization occurred over the average BI baseline in the pre-elevated Mg^2+^ condition. The box-and-whisker plots show normalized BI over a five-minute interval in the 0, 1 mM, 3 mM and 5 mM conditions. Each condition contains up to five cultures with 12 measurements made over the hour-long recording. Red horizontal bars are the median. Black bars stretch to the 25th and 75th percentiles. Red + indicates outliers. All cultures are perfused with the same control medium in the control and washout groups, where in these cases concentration represents their concentration in the drug condition. Black lines show linear regression. Dotted line shows baseline and is consistent with no change in BI relative to control. *A*: With elevated Mg^2+^, there is a significant positive correlation between normalized BI and dose (*p <* 0.05). *B*: In the washout condition, significance is lost (*p >* 0.05).

*Qualitative Observations from Time Rasters* Time rasters were examined to further discern the behavior within cultures. Time rasters are a useful method of showing neuronal firing patterns across a network where time is plotted on the *x*-axis and each neuron is indexed on the *y*-axis. Random portions of time rasters were sampled from each culture in the control, medium change and washout conditions. Time rasters for two representative cultures, one with a sham medium change and one with elevated Mg^2+^, are shown in Figure 3.

**Figure 3.**
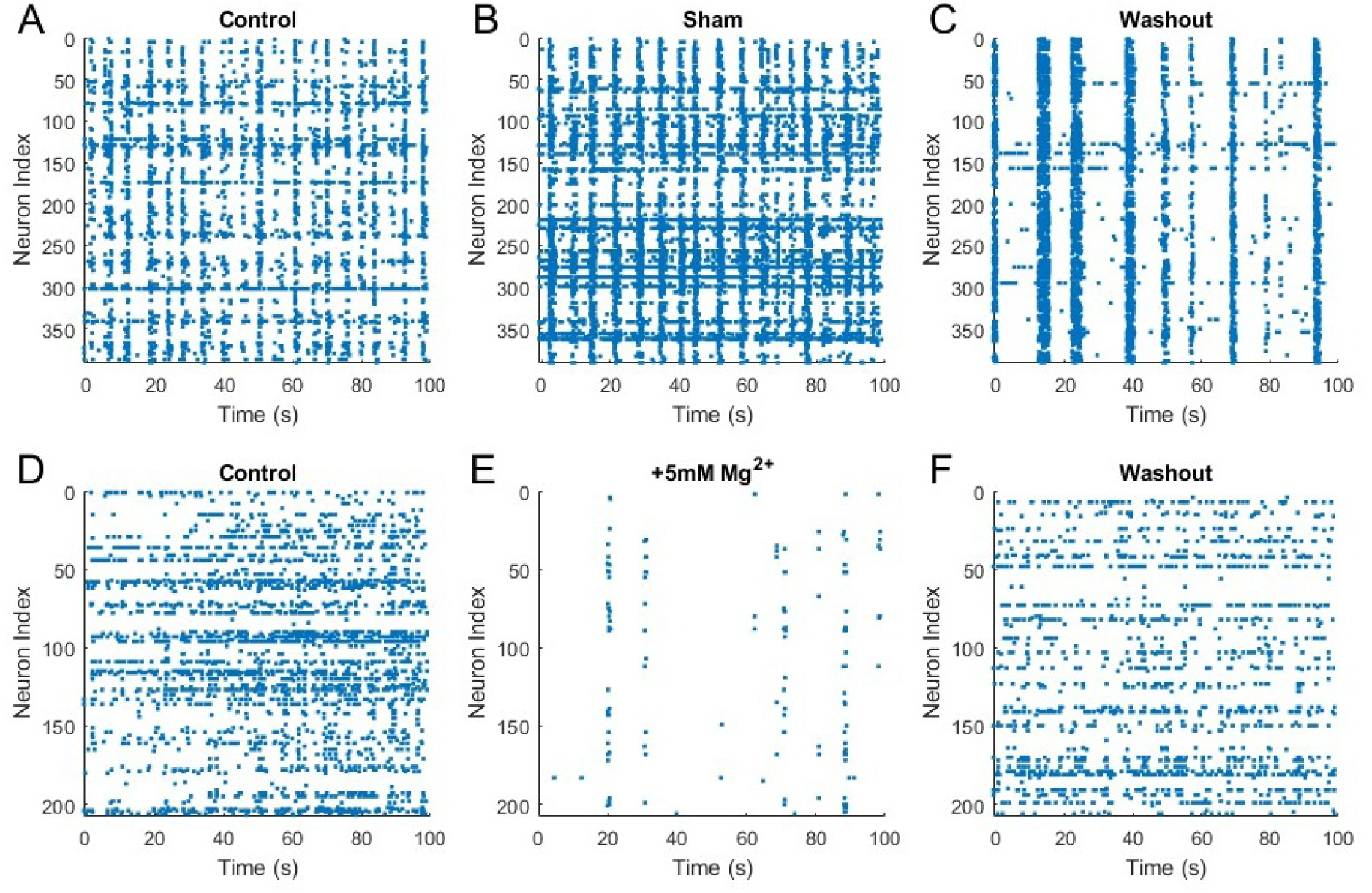
Time rasters for two sample cultures that underwent a sham change of control media (*A-C*) or a 5 mM increase in extracellular Mg^2+^ (*D-F*). *A*: A culture in control medium shows network bursts (vertical stripes), high firing rate neurons (horizontal stripes) and sporadic activity. *B-C*: Over time and with medium changes that do not increase Mg^2+^ concentration, the culture’s activity levels may fluctuate, but all three types of activity are still present. *D*: A different culture in control medium shows primarily tonic spiking and sporadic activity. *E*: When the culture is exposed to medium with 5 mM elevated Mg^2+^, the activity changes considerably from its control condition. All tonically spiking neurons and nearly all sporadic activity vanish, and the only activity present are synchronous bursts. *F*: Washout returns the culture to a state similar to its control condition.

Neuronal cultures show a high degree of heterogeneity, where patterns of neuronal activity can evolve over time or vary even within a different spatial sampling of the same culture Wagenaar, Pine, and Potter (2006a). A simple way to break down this individualized and complex behavior is to consider three activity patterns that are present in time rasters: tonic spiking of high firing-rate neurons (horizontal stripes); synchronous network bursts (vertical stripes); and sporadic spiking (background noise). An increase in network bursting would be represented by an increase in the number of vertical stripes. Since BI calculates the level of bursting relative to other types of activity, an increase in BI would be represented by increased synchronous bursting relative to other types of activity.

Cultures that underwent a sham medium trial (such as Figure 3*A-C*) showed a variety of changes over time. Firing rate changes were noted when changing the culture medium at the beginning of the sham trial and washout, though changes in activity occurred throughout the 3-hour recording window. A given culture may show a gradual increase, decrease, or approximately stable level of activity over time. What remained consistent was that the presence of any of the three types of activity was stable throughout recording. A culture that showed only tonic spiking in the first hour without network bursts or additional sporadic activity would continue to show only tonic spiking.

With elevated levels of Mg^2+^, the time rasters were sparser, showing an overall decrease in activity. This effect appeared to scale with dose, with the greatest effects shown at the highest dose of 5 mM (e.g. Figure 3*E*). 1 mM trials showed the weakest changes. Additional sporadic activity was still present for each trial, and half of cultures showed a significant decrease in the number of synchronous bursts and tonically spiking neurons. Changes were greater in the 3 mM and 5 mM trials. Additional sporadic activity vanished in all but one 3 mM trial and every 5 mM trial. The number of tonically bursting neurons decreased in every 3 mM and 5 mM trial. The changes in synchronous bursts were more complex. Cultures that had a high number of synchronous bursts in the control often showed a slight decrease in elevated Mg^2+^ condition. However, two cultures that showed no synchronous bursts in the control condition developed them with 5 mM elevated Mg^2+^ (e.g. Figure 3*D-F*). In both cases, the synchronous bursting did not remain in the washout condition.

The decrease in network activity caused by elevated Mg^2+^ is clearly shown by the average neuronal firing rates (Figure 4*A*). A significant decrease in firing rate with elevated Mg^2+^ was shown by the Kruskal-Wallis test for the higher doses of 3 mM (*X*^2^ = 5.3333*, p* = 0.0209) and 5 mM (*X*^2^ = 5.7709*, p* = 0.0163). There was no significant firing rate change for the 1 mM dose (*X*^2^ = 0.0833*, p* = 0.7728) or the sham changes (*X*^2^ = 0.0982*, p* = 0.7540). Individual cultures with high doses of Mg^2+^ showed this change in their firing rate (Figure 4*B*) and time rasters (Figure 4*C*). This effect occurred in every culture with 3 mM and 5 mM elevated Mg^2+^ (excluding networks with *<*10 neurons post-spike sorting that did not enable statistical analysis due to undersampling).

**Figure 4.**
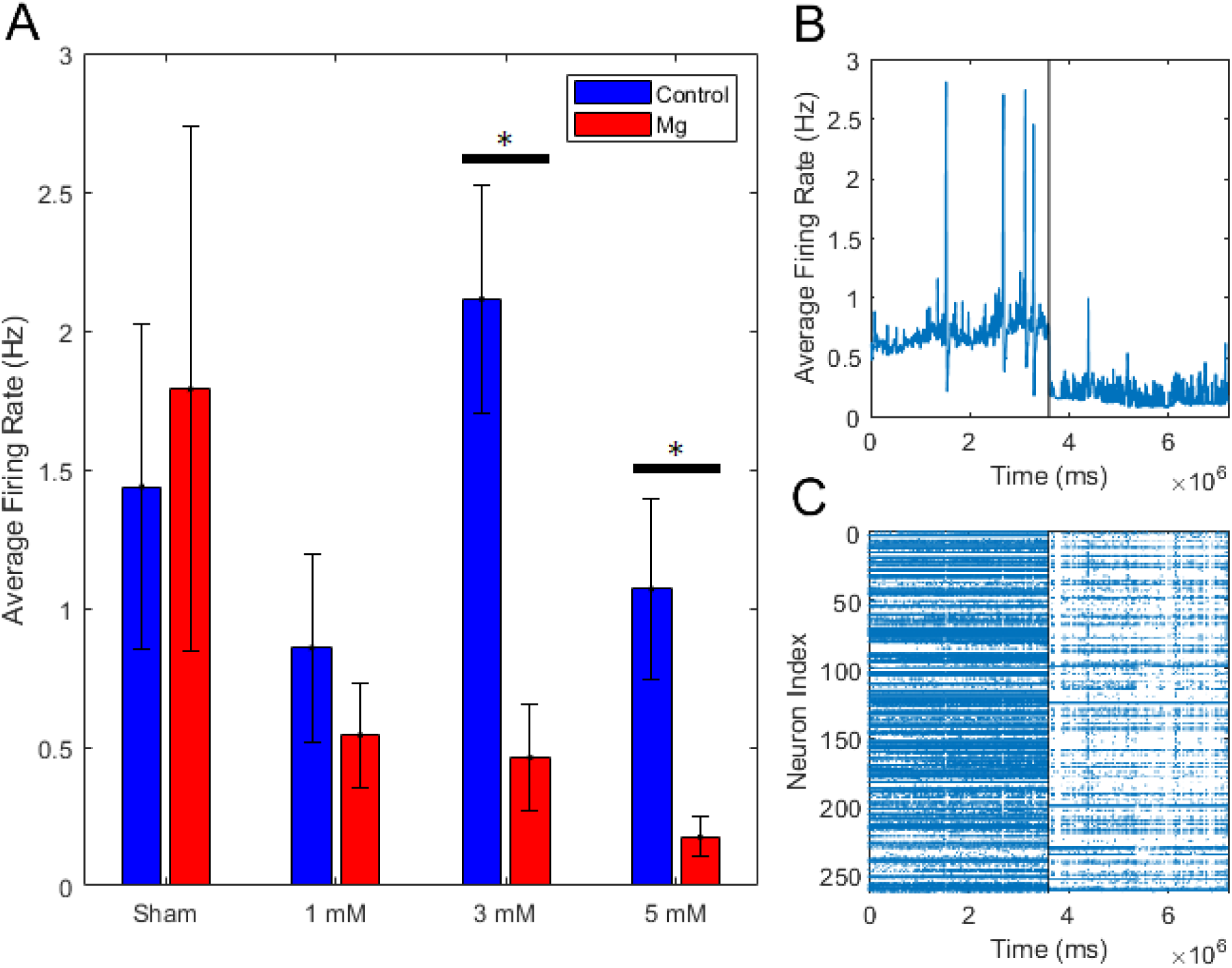
Firing rate drops with elevated magnesium compared to control recordings. *A*: Average firing rate of cultures in control medium (blue) compared to elevated Mg^2+^ (red) for each concentration. Average firing rate did not significantly change for the sham or 1 mM Mg^2+^ changes, but significantly decreased for the 3 mM and 5 mM changes. Error bars represent SEM. *B*: Average firing rate for one representative culture. Black line represents the transition from control medium to medium with 5 mM elevated Mg^2+^, during which the firing rate substantially decreased. *C*: Time raster for the same culture pictured in *B*. Blue dots represent a single spike. In control medium, the spiking frequency is noticeably higher than in medium with 5 mM elevated Mg^2+^.

*Culture Type* Many of the previous studies investigating the effect of Mg^2+^ on neuronal cultures focused on dissociated cultures, e.g. Wagenaar et al. (2006b). Dissociated cultures are often prepared by enzymatically and mechanically dissociating a section of tissue from the brain Hilgenberg and Smith (2007). Due to this process, they are a homogeneous population of neurons lacking a cortical structure. Organotypic slice cultures are prepared from cortical slices of rat pups. As such, they retain some basic features of cortical organization that are not present in dissociated, or primary, cultures. For example, organoytpic cultures typically have distinct cortical layers Bolz, Novak, Götz, and Bonhoeffer (1990); Götz and Bolz (1992), though these layers are not as clear as seen in acute slices and are disrupted Staal, Alexander, Liu, Dickson, and Vickers (2011). Slices from the brain are plated and given a couple weeks to recover prior to recording. Coronally sliced cultures retain all six layers of cortex and a diverse population of neurons.

To discern whether the bursting results are specific to one type of culture preparation, BI was compared across different culture types, including the neocortex of organotypic cultures recorded here, dissociated hippocampal cultures Timme et al. (2016) and dissociated neocortical cultures Wagenaar et al. (2005). Cultures were not controlled for number or density of neurons, which can have a significant effect upon burstiness Wagenaar et al. (2006b).

The same average BI was found for control recordings of organotypic (0.23 ± 0.035) and dissociated hippocampal cultures (0.240 ± 0.014*, p* = 0.68274). However, dissociated neocortical cultures (0.48 ± 0.02) showed a significantly higher BI than both organotypic neocortical(*p* = 1.7335 ∗ 10^−7^) and dissociated hippocampal cultures (*p* = 1.7737 ∗ 10^−7^).

### Information Dynamics

Following network-wide analysis, the information dynamics of individual or small groups of neurons were considered. The information theoretic measures are broken into first-order measures, which involve individual neurons, and higher-order measures, which involve pairs or triads of neurons. When comparing the elevated Mg^2+^ condition to control and washout, all doses (1, 3, 5 mM) were pooled into a single elevated Mg^2+^ condition. Pooling the data provided the largest possible sample size and avoided biasing the data given no *a priori* knowledge of dose responses. Individual doses were not directly compared to the control and washout conditions, but relationships between them are shown.

*First Order Measures* The Friedman’s *X*^2^ test showed a significant effect on Shannon entropy between conditions (Figure 5*A*, *X*^2^ = 319.4972*, p* = 4.1885 ∗ 10^−70^). A Wilcoxon post-hoc test showed that entropy was lowest in the elevated Mg^2+^ condition ((6.19 ± 0.31) ∗ 10^−3^), significantly lower than both the control ((1.370 ± 0.053) ∗ 10^−2^*, p* = 1.4716 ∗ 10^−86^) and washout ((1.045 ± 0.042) ∗ 10^−2^*, p* = 2.8864 ∗ 10^−31^). The washout was significantly lower than the control condition (*p* = 3.4016 ∗ 10^−14^), showing that it did not fully return to its baseline after an hour. The Kruskal-Wallis test showed that entropy also had a highly significant dose-dependent effect (Figure 5*B*, *X*^2^ = 456.9112*, p* = 6.0642 ∗ 10^−100^) and decreased with higher concentrations. Post-hoc testing using the Mann-Whitney U-test showed that the 1 mM dose ((9.45 ± 0.52) ∗ 10^−3^) had significantly higher entropy than both 3 mM ((6.23 ± 0.58) ∗ 10^−3^*, p* = 2.6214 ∗ 10 − 26) and 5 mM ((2.34 ± 0.35) ∗ 10^−3^*, p* = 1.787 ∗ 10^−96^). The 3 mM dose had significantly higher entropy than 5 mM (*p* = 1.8641 ∗ 10^−32^).

**Figure 5.**
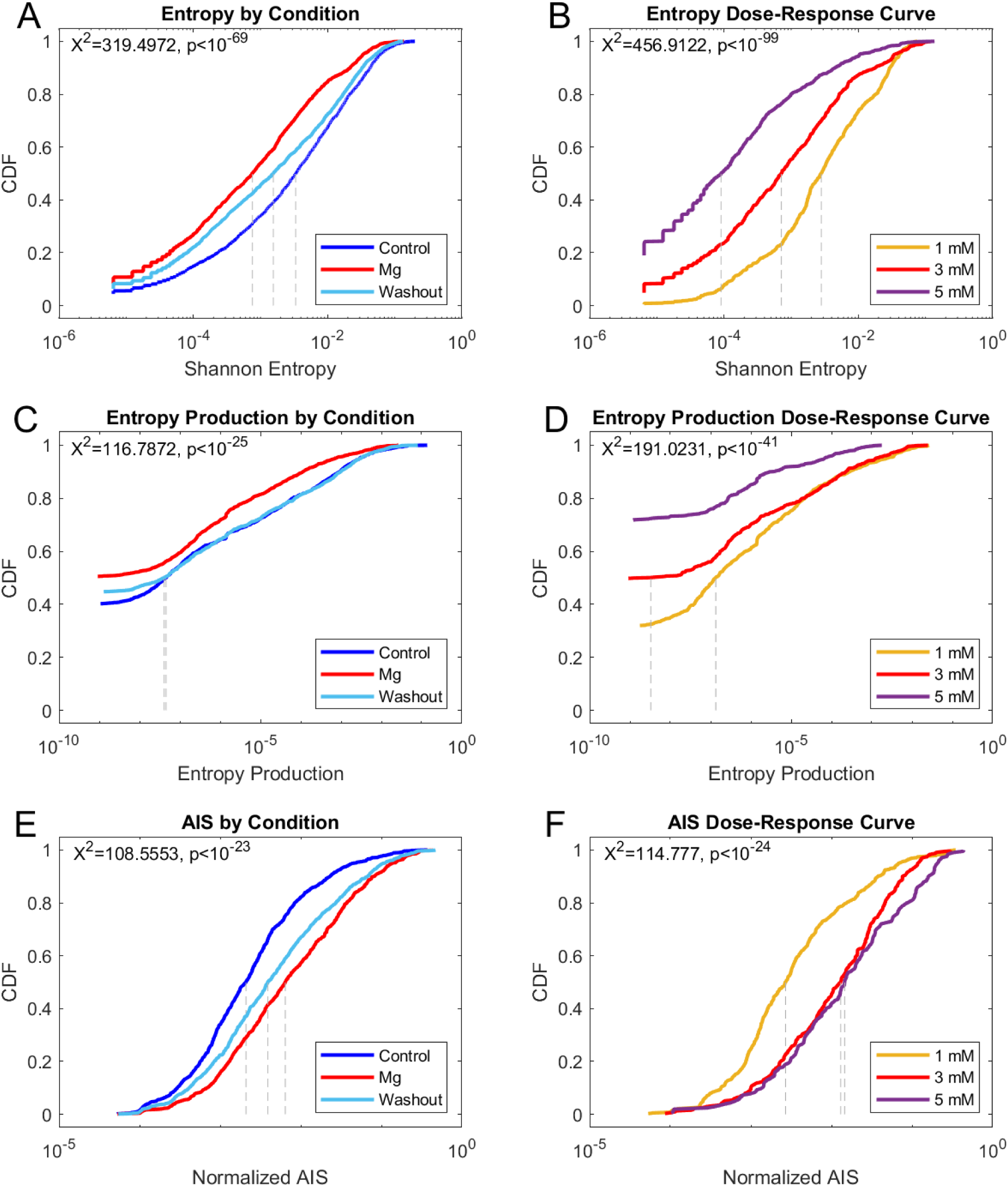
Cumulative distribution function (CDF) of first-order information dynamic measures by recording condition and dose of Mg^2+^. Dotted gray lines represent median values. Significant effects by recording condition and dose were found in all measures. *A*: CDF for Shannon entropy for each recording condition (dark blue: control, red: elevated Mg^2+^ for all doses, light blue: washout) *B*: CDF for Shannon entropy for each dose of Mg^2+^ (yellow: 1 mM, red: 3 mM, purple: 5 mM). *C-F*: CDFs given by condition and dose for (*C-D*) entropy production and (*E-F*) active information storage (AIS).

Friedman’s *X*^2^ showed a small but significant effect by condition on entropy production (Figure 5*C*, *X*^2^ = 116.7872*, p* = 4.365 ∗ 10^−26^). The elevated Mg^2+^ condition ((2.71 ± 0.36) ∗ 10^−4^) had significantly lower entropy production than both the control ((8.3 ± 1.1) ∗ 10^−4^*, p* = 8.631 ∗ 10^−25^) and washout ((6.74 ± 0.76) ∗ 10^−4^*, p* = 3.187 ∗ 10^−30^). In this measure, the washout completely removed the effect, as the control and washout showed no significant difference (*p* = 0.60013). Kruskal-Wallis showed that the dose of Mg^2+^ also had a significant effect on entropy production (Figure 5*D*, *X*^2^ = 191.0231*, p* = 3.3103 ∗ 10^−42^). Once again, entropy production was significantly higher for the 1 mM dose ((4.51 + 0.83) ∗ 10^−4^) than both 3 mM ((3.00 ± 0.57) ∗ 10^−4^*, p* = 2.1997 ∗ 10^−6^) and 5 mM ((1.98 ± 0.45) ∗ 10^−5^*, p* = 1.3508 ∗ 10^−44^), and significantly higher for 3 mM than 5 mM (*p* = 2.2602 ∗ 10^−18^).

Friedman’s *X*^2^ also showed a significant effect of AIS by condition (Figure 5E, *X*^2^ = 108.5553*, p* = 2.6762 ∗ 10^−24^). AIS was significantly higher with elevated Mg^2+^ (0.0350 ± 0.0029) than both the control (0.0144 ± 0.0018*, p* = 8.0531 ∗ 10^−31^) and washout (0.0295 ± 0.0029*, p* = 2.3381 ∗ 10^−4^). Again, there was not a complete recovery in the washout condition, as it was significantly higher than the control (*p* = 4.3724 ∗ 10^−16^). The Kruskal-Wallis test showed significance in the dose-response, as well (Figure 5F, *X*^2^ = 114.777*, p* = 1.1926 ∗ 10^−25^). The 1 mM dose (0.0166 ± 0.0023) had significantly lower AIS than both 3 mM (0.0299 ± 0.0025*, p* = 6.8361 ∗ 10^−19^) and 5 mM (0.0498 ± 0.0053*, p* = 3.2554 ∗ 10^−19^). There was no difference between 3 mM and 5 mM (*p* = 0.092107), suggesting that this effect was saturated at higher doses.

Taken together, the first-order measures showed that elevated Mg^2+^ decreases Shannon entropy and entropy production and increases AIS. These effects were apparent by condition and generally increased with concentration of Mg^2+^. All of these effects suggest a decrease in the complexity of the neuronal activity with elevated Mg^2+^. The network had more time-reversible, predictable dynamics that did not vary significantly.

*Second- and Higher-Order Measures* Friedman’s *X*^2^ showed a significant effect of mTE by condition (Figure 6*A*, *X*^2^ = 607.3753*, p* = 1.2886 ∗ 10^−132^). A schematic representation of mTE is shown in Figure 6 where each line represents a significant connection. Sample connectivity matrices are included in Supplemental Figures 1 and 2. Similar to several previous measures, mTE was greatest in the elevated Mg^2+^ condition ((2.87 ± 0.13) ∗ 10^−2^), significantly higher than both the control ((1.73 ± 0.12) ∗ 10^−2^*, p* = 4.1889 ∗ 10^−85^) and washout ((1.97 ± 0.11) ∗ 10^−2^*, p* = 9.5132 ∗ 10^−70^). This effect was not completely reversed with washout, which had significantly greater mTE than the control (*p* = 5.4975 ∗ 10^−18^). The dose-response also showed significance based upon the Kruskal-Wallis test (Figure 6*B*, *X*^2^ = 566.6805*, p* = 8.8488 ∗ 10^−124^). The 1 mM dose ((1.902 ± 0.090) ∗ 10^−2^) had significantly lower mTE than 3 mM ((2.439 ± 0.050) ∗ 10^−2^*, p* = 2.5618 ∗ 10^−92^) and 5 mM ((3.100 ± 0.087) ∗ 10^−2^*, p* = 1.6071 ∗ 10^−87^). 3 mM was in the middle, with significantly lower mTE than 5 mM (*p* = 1.0238 ∗ 10^−15^).

**Figure 6.**
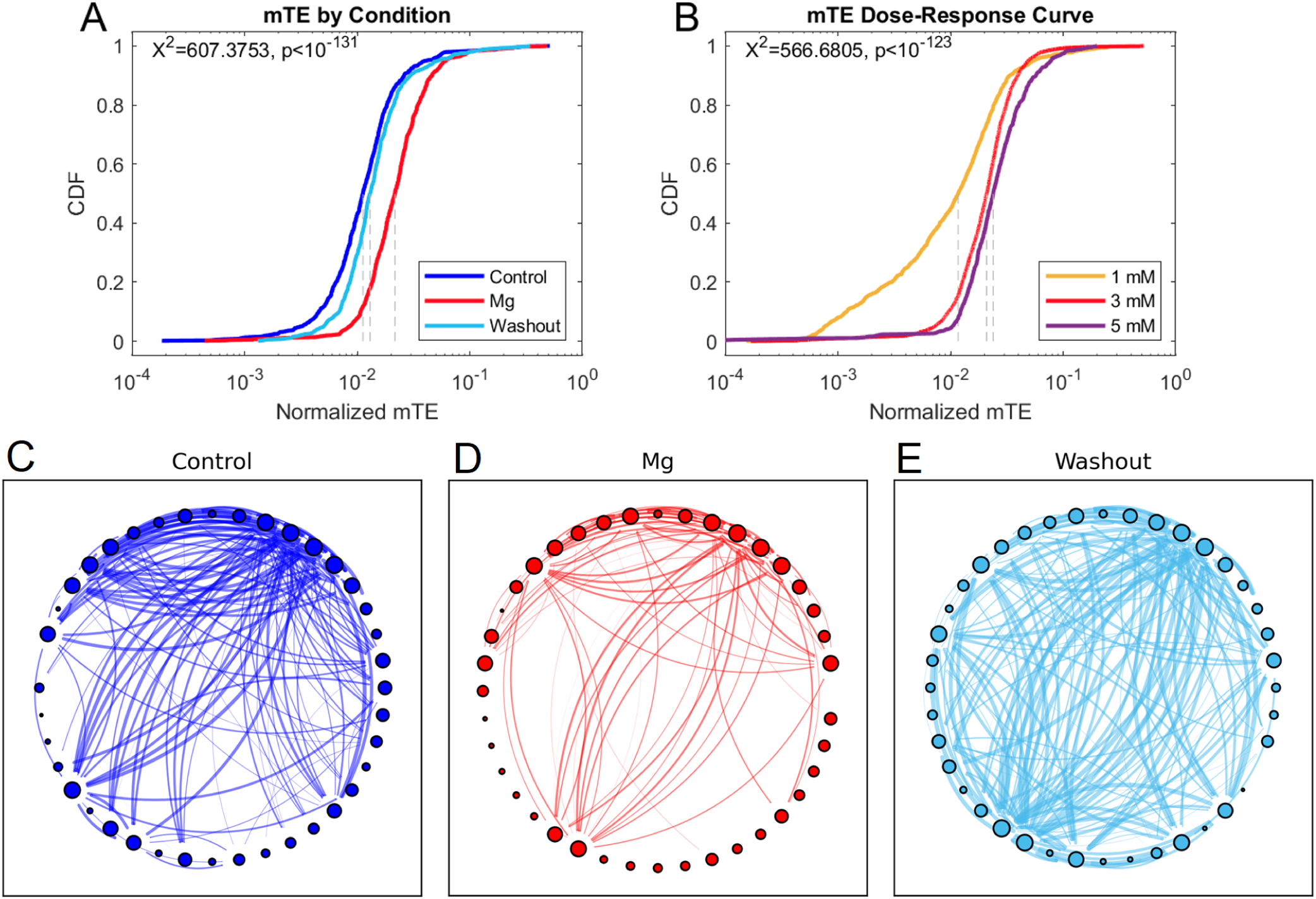
Cumulative distribution function (CDF) and network diagrams of multivariate transfer entropy (mTE). *A*: CDF for mTE for each recording condition (dark blue: control, red: elevated Mg^2+^ for all doses, light blue: washout). Dotted gray lines represent median values. Significant effects by recording condition and dose were found in all measures. *B*: CDF for mTE for each dose of Mg^2+^ (yellow: 1 mM, red: 3 mM, purple: 5 mM). *C*: Network structure of a representative control network showing significant connections as calculated by mTE. Each circle represents a neuron, where the size scales with the degree of the neuron. Strength of the connection given by mTE is shown by the thickness of the line. The same network is shown during *(D)* the elevated Mg^2+^ condition and *(E)* the washout condition.

In addition to overall mTE, Friedman’s *X*^2^ showed significant effects for distributions of in-degree (*X*^2^ = 145.9766*, p* = 2.0026 ∗ 10^−32^) and out-degree of neurons (*X*^2^ = 72.5984*, p* = 1.7197 ∗ 10^−16^). These measures represent the number of significant incoming and outgoing connections for neurons, respectively. The in-degree had significantly more edges for control (3.537 ± 0.089) than both the elevated Mg^2+^ (2.368 ± 0.078*, p* = 1.9629 ∗ 10^−36^) and washout (3.190 ± 0.094*, p* = 8.4633 ∗ 10^−4^).

Mg^2+^ had a significantly lower in-degree than washout (*p* = 2.151 ∗ 10^−18^). Out-degree similarly showed that the control condition (3.59 ± 0.24) had significantly more edges than elevated Mg^2+^ (2.37 ± 0.20*, p* = 4.3561 ∗ 10^−14^) and washout (3.24 ± 0.24*, p* = 1.8866 ∗ 10^−3^). In this case, the washout condition also had significantly more edges than the elevated Mg^2+^ condition (*p* = 3.9999 ∗ 10^−5^). These effects of degree are visible in Figure 6, where the elevated Mg^2+^ condition is shown to have significantly fewer highly-connected neurons than the control or washout. The degree distributions were also significant by dose for both in-degree (*X*^2^ = 105.4088*, p* = 1.2905 ∗ 10^−23^) and out-degree (*X*^2^ = 35.1673*, p* = 2.3095 ∗ 10^−8^). The in-degree was highest for the 3 mM dose (3.74 ± 0.15), significantly above both the 1 mM (1.8449 ± 0.097*, p* = 1.9562 ∗ 10^−14^) and 5 mM dose (1.63 ± 0.12*, p* = 2.7131 ∗ 10^−20^). 1 mM had the next highest in-degree, which was significantly higher than 5 mM (*p* = 2.1557 ∗ 10^−4^). The results for out-degree were similar. The 3 mM dose produced significantly higher out-degree (3.74 ± 0.43) than 5 mM (1.63 ± 0.27*, p* = 7.6008 ∗ 10^−7^) but not 1 mM (1.84 ± 0.21*, p* = 0.56184). 1 mM had significantly higher out-degree than the 5 mM dose (*p* = 5.9999 ∗ 10^−9^).

Clustering coefficient had a small but significant effect according to Friedman’s *X*^2^ (Figure 7*A*, *X*^2^ = 6.2781*, p* = 0.043324). The elevated Mg^2+^ had the highest clustering coefficient (0.659 ± 0.014) compared to control (0.623 ± 0.013*, p* = 7.8116 ∗ 10^−3^) and washout (0.633 ± 0.014*, p* = 0.022993). The clustering coefficient for control and washout showed no statistical difference (*p* = 0.24375). There was also a significant effect of dose given by the Kruskal-Wallis test (Figure 7*B*, *X*^2^ = 28.9478*, p* = 5.1767 ∗ 10^−7^). Clustering coefficient was significantly higher for the 5 mM dose (0.785 ± 0.022) than 1 mM (0.643 ± 0.019*, p* = 4.3955 ∗ 10^−6^) or 3 mM (0.692 ± 0.012*, p* = 2.5338 ∗ 10^−7^). The 1 mM and 3 mM doses did not show a significant difference in clustering coefficient (*p* = 0.28139).

**Figure 7.**
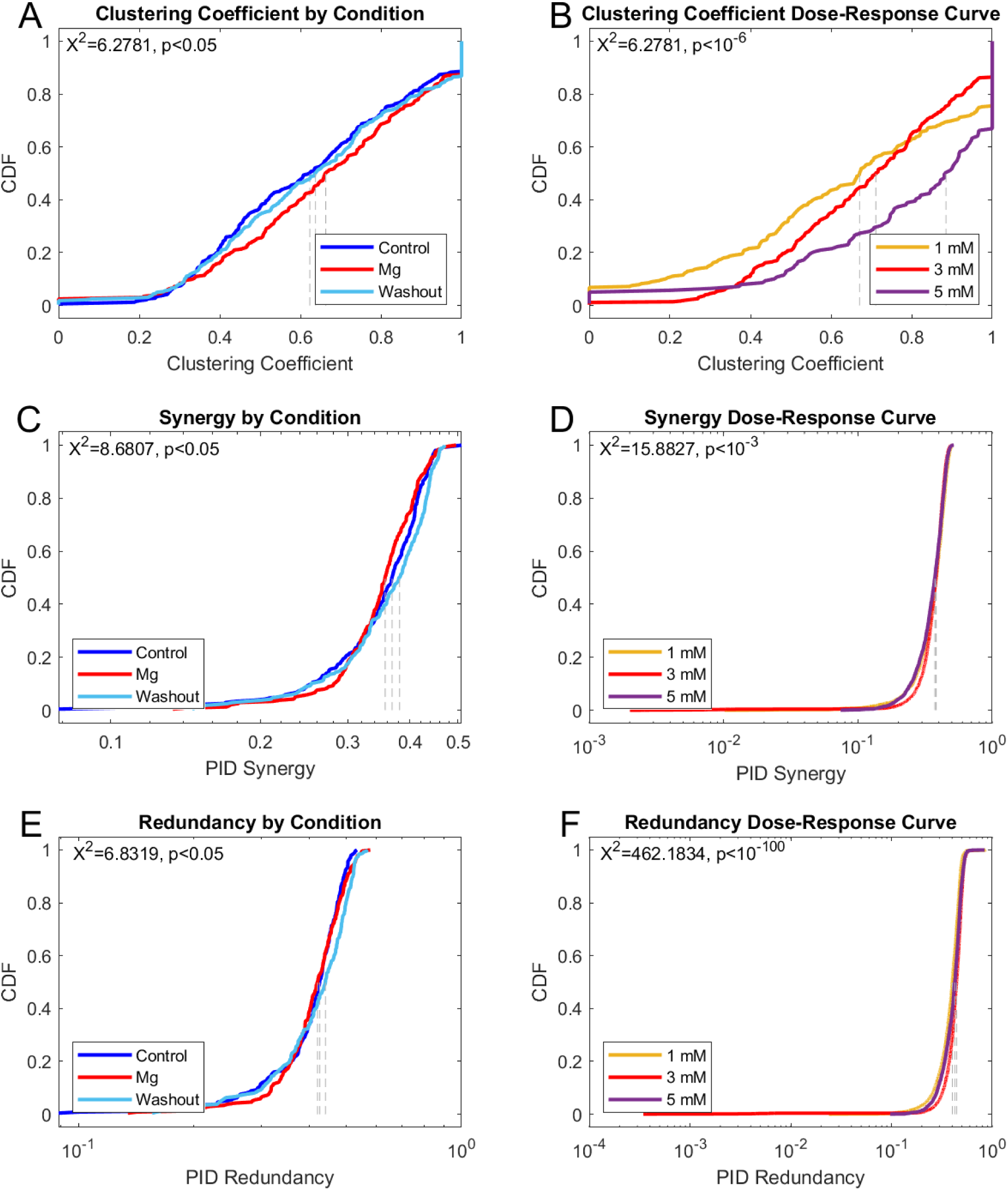
Cumulative distribution function (CDF) of second- and higher-order information dynamic measures by recording condition and dose of Mg^2+^. Dotted gray lines represent median values. Significant effects by recording condition and dose were found in all measures. *A*: CDF for clustering coefficient for each recording condition (dark blue: control, red: elevated Mg^2+^ for all doses, light blue: washout). *B*: CDF for clustering coefficient for each dose of Mg^2+^ (yellow: 1 mM, red: 3 mM, purple: 5 mM). *C-F*: CDFs given by condition and dose for (*C-D*) synergy and (*E-F*) redundancy computed from partial information decomposition (PID).

Friedman’s *X*^2^ showed that synergy had a small but significant effect by condition (Figure 7*C*, *X*^2^ = 8.6807*, p* = 0.013032). Wilcoxon pairwise post-hoc testing revealed that there was no significant difference in synergy between elevated Mg^2+^ (0.3530 ± 0.0041) and the control (0.3548 ± 0.0047*, p* = 0.62181). However, elevated Mg^2+^ had significantly higher synergy than the washout (0.3647 ± 0.0050*, p* = 0.0070986) and control had significantly lower synergy than the washout (*p* = 0.023157). Synergy also had a significant overall dose-dependent response (Figure 7*D*, *X*^2^ = 15.8827*, p* = 3.5572 ∗ 10^−4^). The 5 mM dose (0.3607 ± 0.0019) had significantly lower synergy than both 1 mM (0.3674 ± 0.0018*, p* = 4.6536 ∗ 10^−4^) and 3 mM (0.37082 ± 0.00088*, p* = 2.9819 ∗ 10^−4^). There was no significant difference between synergy in the 1 mM and 3 mM doses (*p* = 0.31407).

Friedman’s *X*^2^ test showed a significant effect by condition on redundancy (7*E*, *X*^2^ = 6.8319*, p* = 0.032845). However, the Wilcoxon paired post-hoc tests showed that there was no significant difference between elevated Mg^2+^ (0.4154 ± 0.0049) and control (0.4096 ± 0.0053, *p* = 0.33622) or elevated Mg^2+^ and washout (0.4225 ± 0.0058*, p* = 0.16753). Strangely, significance occurred only between the control and washout (*p* = 0.0039378) with the control having significantly lower redundancy. Dose also had a significant overall effect on redundancy (7*F*, *X*^2^ = 462.1834*, p* = 4.3467 ∗ 10^−101^). The 1 mM dose (0.3884 ± 0.0018) had significantly lower redundancy than the 3 mM (0.4308 ± 0.0010*, p* = 2.5901 ∗ 10^−100^) and 5 mM doses (0.4108 ± 0.0020*, p* = 2.0517 ∗ 10^−17^). Interestingly, redundancy was greatest for the 3 mM dose and was significantly higher than 5 mM (*p* = 4.4063 ∗ 10^−19^).

The higher-order measures show an increased consistency in information transfer. The significant increase of mTE with elevated Mg^2+^ indicates raised network connectivity, while increased redundancy shows that neurons are transmitting the same information at a greater rate. The weak results in synergy suggests that the network is more cohesive but may perform fewer computations.

**Table 1.** Summary of information dynamic measures. All doses of elevated Mg^2+^ (1, 3, 5 mM) are compared to the control and washout conditions. The three doses are separately considered. Comparisons represent statistically significant differences found between the groups.

| Measure | By Condition | By Dose |
| --- | --- | --- |
| Entropy | Mg < washout < control | 5mM < 3mM < 1mM |
| Entropy Production | Mg < (control = washout) | 5mM < 1mM < 3mM |
| AIS | control < washout < Mg | 1mM < 3mM < 5mM |
| mTE | control < washout < Mg | 1mM < 3mM < 5mM |
| In-Degree | Mg < washout < control | 1mM < 5mM < 3mM |
| Out-Degree | Mg < washout < control | 5mM < (1mM = 3mM) |
| Clustering Coefficient | (control = washout) < Mg | (1mM = 3mM) < 5mM |
| Synergy | (control = Mg) < washout | 5mM < (3mM = 1mM) |
| Redundancy | control = Mg, washout = Mg,<br>control < washout | 1mM < 5mM < 3mM |

## DISCUSSION

The burstiness index (BI) showed a significant dose-dependent increase with elevated Mg^2+^ compared to control and washout conditions. Observations of the time rasters suggested that the increase in BI with elevated Mg^2+^ was not caused by a simple increase in the burst rate. Two cultures showed an emergence of bursting with 5 mM Mg^2+^ that was not present in the control conditions or washout. Other cultures with elevated Mg^2+^ appeared to show a decrease in the number of high firing rate neurons and a substantial decrease in sporadic activity outside of bursts or tonically spiking neurons. Both cases resulted in an increase in the ratio of network bursting to other types of activity.

Information dynamics painted a deeper picture. First-order measures (entropy, entropy production and AIS) showed that increasing Mg^2+^ reduces network activity. The remaining activity is more time-reversible with a more predictable future state. Higher-order measures (mTe, in- and out-degree, clustering coefficient, synergy and redundancy) showed that overall network connectivity and clustering are increased, but with weaker connections. There is a heightened transfer of redundant information.

Synergistic information shows a slight decrease, suggesting that less computation is occurring within the network. The effects of elevated Mg^2+^ seem to be two-fold: they raise the threshold for neuron firing; and when neurons do fire, the network is highly integrated where clustered neurons fire collectively in a predictable and redundant manner.

In a majority of information theoretic measures, the washout condition yielded results between the control and elevated Mg^2+^ conditions, indicating that an hour-long washout with control medium did not fully return the network to its baseline values.

Contrary to the other information theoretic measures, the redundancy, entropy production and in-degree were larger in the 3 mM dose than 1 mM or 5 mM. There is a chance this is caused by the small effect size or a systematic difference in the 3 mM group of cultures compared to 5 mM. Otherwise, it is possible that this is an effect that saturates at 3 mM. Substances applied to neuronal cultures have previously been shown to have network activity metrics that are greatest at a particular dose Stewart and Plenz (2006). High levels of Mg^2+^ have noted cytotoxic effects on cultures Baker et al. (1991), and 5 mM exceeds values typically required for burst quieting, e.g. Maeda et al. (1995). 3 mM may be the dose that maximizes redundancy, entropy production and in-degree. This saturation effect may have also played a role in the 3 mM group not showing a decreased BI during washout.

Our results suggest that the canonical view of Mg^2+^ as a “burst quieter” is limited. Previously, it was shown that burstiness decreases with 1-2 mM elevated extracellular Mg^2+^ Wagenaar et al. (2006b). This is not necessarily at odds with our increase in BI, since BI shows an increase in the *ratio* of bursts relative to other types of activity. However, some cultures showed an appearance of bursting with Mg^2+^ that was not present in the control or washout.

A potential explanation for the increase in burst rate is that culture type impacted the interaction of Mg^2+^. We recorded the cortex of organotypic cultures, which somewhat retain cortical layers, while many previous burstiness studies used dissociated cultures of various populations, e.g. Wagenaar et al. (2005). We found that the spontaneous burst rate of organotypic cultures was statistically the same as dissociated hippocampal cultures but significantly lower than dissociated neocortical cultures. This effect may be caused by differences in culture size or density, which can significantly impact burstiness Wagenaar et al. (2006a). We expect that a smaller culture size and high neuronal density result in high burstiness Timofeev, Grenier, Bazhenov, Sejnowski, and Steriade (2000); Wagenaar et al. (2006a) due to increased neuronal connectivity. Our estimates suggest that the sizes and densities of these cultures increased proportionally: size and density were smallest for the neocortical dissociated cultures, approximately twice as high for the hippocampal dissociated cultures, and approximately four times higher still for the neocortical organotypic cultures Korbo et al. (1990); Stoppini, Buchs, and Muller (1991); Su, Paradiso, Long, Liao, and Simonato (2011); Tang et al. (2008). The proportional increase of both size and density cancel any expectations of changes in burstiness. This may account for the similarity in organotypic neocortical and dissociated hippocampal culture BI, but does not clearly explain why dissociated neocortical cultures have a significantly lower BI. It would be interesting to make such a comparison in an experiment that controls for culture size and density across different culture preparations, comparing organotypic and dissociated cultures taken from cortex and hippocampus. Each dose of Mg^2+^ resulted in one organotypic culture with an average BI that decreased rather than increased due to biological variation within samples. Further, we observed that having a high or low average control BI did not predict whether average BI with elevated Mg^2+^ would increase or by how much (data not shown). Taken together, this indicates that spontaneous BI does not predict an increase in BI with Mg^2+^.

A stronger explanation is that the layered structure of the cortex results in complex dynamics that are inhibited by elevated Mg^2+^. NMDAR are most numerous in layers II/III and are a significant mediator of sending signals downstream to layer V Currie et al. (1994). Complex dynamics such as neuronal avalanches Beggs and Plenz (2003) primarily occur in layers II/III. A decrease in these types of complex behavior is consistent with our findings that Mg^2+^ leads to highly integrated and predictable network activity, and one with few spikes not contained in a synchronous burst or a tonically spiking neuron. With complex dynamics inhibited, the only network activity that may be able to surpass the Mg^2+^ block is a network-wide burst. This is similar to using Mg^2+^ in stimulation studies to block all activity except bursts caused by electrical stimulation, e.g. Wagenaar et al. (2006b). While this study focused on the dynamics of Mg^2+^ between individual, pairs and triplets of neurons across the entire network, using post-hoc staining to identify the cortical layers in which those neurons resided would help to elucidate this mechanism. This investigation would be further strengthened by incorporating electrical stimulation to directly test the ways in which Mg^2+^ alters network activity, e.g. Liu, Seay, and Buonomano (2023); Maeda et al. (1998); Shahaf et al. (2008); Wagenaar et al. (2006b).

Network bursts contain a high degree of spatiotemporal complexity Corner et al. (2002). Further suites of analysis could be applied to the existing data, such as a deeper examination of burst patterns by sorting the time rasters by Pearson correlations or connective communities (e.g. Louvain clustering).

Computation of metrics such as population activity Estévez-Priego et al. (2023) would reveal whether Mg^2+^ causes a change in which neurons participate in bursts. Additional experimentation would allow for an even deeper dive into the spatiotemporal dynamics. For example, spatial patterning of bursts could be investigated by constraining culture growth to topographical patterns, e.g. Montalà-Flaquer et al. (2022). Such analyses would synergize well with our information dynamical analysis to further elucidate the impact of Mg^2+^ on bursting.

A further test to determine the impact of NMDAR on the information dynamics would be to target these receptors directly. Elevated Mg^2+^ reduces sporadic activity and burst frequency by raising voltage thresholds Dribben, Eisenman, and Mennerick (2010) and blocking NMDARs. This extends inter-burst intervals for synaptic recovery Tabak et al. (2010), enhancing burst onset recruitment and neuronal synchronization. A similar set of experiments to this work could be run using 2-amino-5-phosphonovaleric acid (APV), an NMDAR antagonist. Elevated Mg^2+^ and APV both canonically reduce bursting by targeting NMDAR. In zero-Mg^2+^ conditions, APV has been shown to stop dissociated cultures from spontaneously bursting Robinson et al. (1993) and to attenuate bursts in hippocampal cultures Mangan and Kapur (2004). In organotypic cultures, APV causes a strong acute decrease in firing and burst rate and a prolonged effect on network bursting if left on chronically Corner et al. (2002). Future experiments applying APV to neuronal cultures would serve as an ideal control for testing the increase in BI. It would be interesting to observe whether APV causes any differences in information dynamics relative to elevated Mg^2+^.

Information dynamics were previously used to investigate the effects of the psychedelic N,N-dipropyltryptamine (DPT) on organotypic slice cultures Varley et al. (2024). Several information dynamical measures show effects opposite of elevated Mg^2+^. DPT showed an increase in entropy and entropy production compared to controls, and a decrease in overall mTE despite an increase in in-degree. Overall, this suggested an increase in the number of weak connections throughout the network. This contrasts with elevated Mg^2+^, which showed a decrease in entropy, entropy production and in-degree, but an increase in mTE. In this context, Mg^2+^ may act akin to an “anti-psychedelic” when the threshold is exceeded in that it increases propagation of highly-correlated and time-reversible activity across the network. However, when the threshold is not exceeded, the network behaves as if it is anesthetized. One similarity is an apparent reduction in synergy: DPT decreased synergy compared to control and washout, while Mg^2+^ had slightly lower synergy than the washout. This suggests the possibility that both psychedelics and Mg^2+^ may inhibit computation in the brain, though the mechanisms that do so differ.

Interestingly, both elevated Mg^2+^ Dolati et al. (2020) and psychedelics Henderson et al. (2025) have been proposed as treatments for migraine, though there is far less clinical data on use of psychedelics for such a purpose. Psychedelics are thought to potentially disrupt cortical spreading depression (CSD), an excessive neuronal excitation followed by depression that spreads across the cortex which is associated with migraine auras preceding the onset of pain. Psychedelics and Mg^2+^ may both disrupt this wave via different mechanisms. Psychedelics may counteract this effect by continuing to propagate activity through its enhanced network of weak connections, thus resisting inhibition. Increasing Mg^2+^ has an overall quieting effect on the network but an increase in redundant and time-reversible activity, both of which may counteract a network-wide inhibitory effect. Further work would need to be done to directly relate these mechanisms to CSD or other features of migraines.

This study focused on acute effects of Mg^2+^ occurring within one hour. Chronic effects five hours after elevated Mg^2+^ application show different effects Slutsky et al. (2010). Cultures exposed to a chronic increase of 0.4 mM Mg^2+^ seemed to homeostatically recover their NMDAR current. Chronic Mg^2+^ blocked NMDAR, resulting in fewer channels opening, but the cultures compensated by releasing more neurotransmitter each time they did. This resulted in an enhancement of NMDA current during network-wide bursts. This may lead to very different information dynamics in cultures treated with chronic rather than acute elevated Mg^2+^. Studying these chronic effects would help to further understand how Mg^2+^ could be used as a treatment for neurological disorders.

We showed that acute elevated Mg^2+^ creates an increased proportion of bursting to other types of neuronal activity in organotypic slice cultures. The network activity becomes highly integrated, showing decreased entropy and increased time-reversibility and connectivity. This is reflected in a loss of neuronal activity not contained within a network burst or a single high firing rate neuron. These results show a loss of complexity that may be tied to disruption of activity within specific cortical layers. This rich network-level analysis on Mg^2+^ in the cortex shows that its effect is more complex than a simple burst quieter. Further comprehension of this simple molecule may be the key to unlocking its potential as a treatment for neurological disorders.

## ACKNOWLEDGMENTS

The authors acknowledge the Indiana University Pervasive Technology Institute for providing supercomputing resources that have contributed to the research results reported within this paper. This research was supported in part by Lilly Endowment, Inc., through its support for the Indiana University Pervasive Technology Institute. This work was supported in part by Shared University Research grants from IBM, Inc., to Indiana University. Additionally, the authors would like to thank Daniel Havert for suggesting an investigation of elevated magnesium on organotypic slices and Naruepon Weerawongphrom for performing dissections.

## TECHNICAL TERMS

**N-methyl-D-aspartate Receptors (NMDAR):** Glutamate receptor often found on pyramidal neurons. Blocked by magnesium ions.

**Burstiness Index (BI):** Dimensionless measure of synchronous neuronal firing within a network relative to total firing.

**Organotypic Culture:** In vitro cortical culture preparation that approximately retains cortical layers.

**Entropy:** High entropy networks visit their states roughly equally (homogeneous, disordered); low entropy networks visit only a few states (inhomogeneous, ordered).

**Entropy Production:** Measure of time reversibility of a process based upon a measured increase in entropy over time.

**Active Information Storage (AIS):** Amount of information that the past state of a neuron reveals about its future state.

**Transfer Entropy:** Information that the past of one or multiple neurons give about the future state of another neuron, beyond its own auto-prediction.

**Degree:** Number of significant connections shared between a pair of neurons.

**Clustering Coefficient:** Measure of how interconnected a neuron’s neighbors are to each other.

**Partial Information Decomposition (PID):** Breakdown of information conveyed by a triad of neurons into synergistic, redundant and unique information passed between them.

**Synergy:** Novel information generated by the convergence of two connected neurons to a single neuron.

**Redundancy:** Information sent twice to a neuron from each of two connected neurons.

