## Supplementary figures and images for "Effect of acute elevated magnesium on bursting activity and information-processing dynamics in cortical cultures"

### Supplemental Figure 1

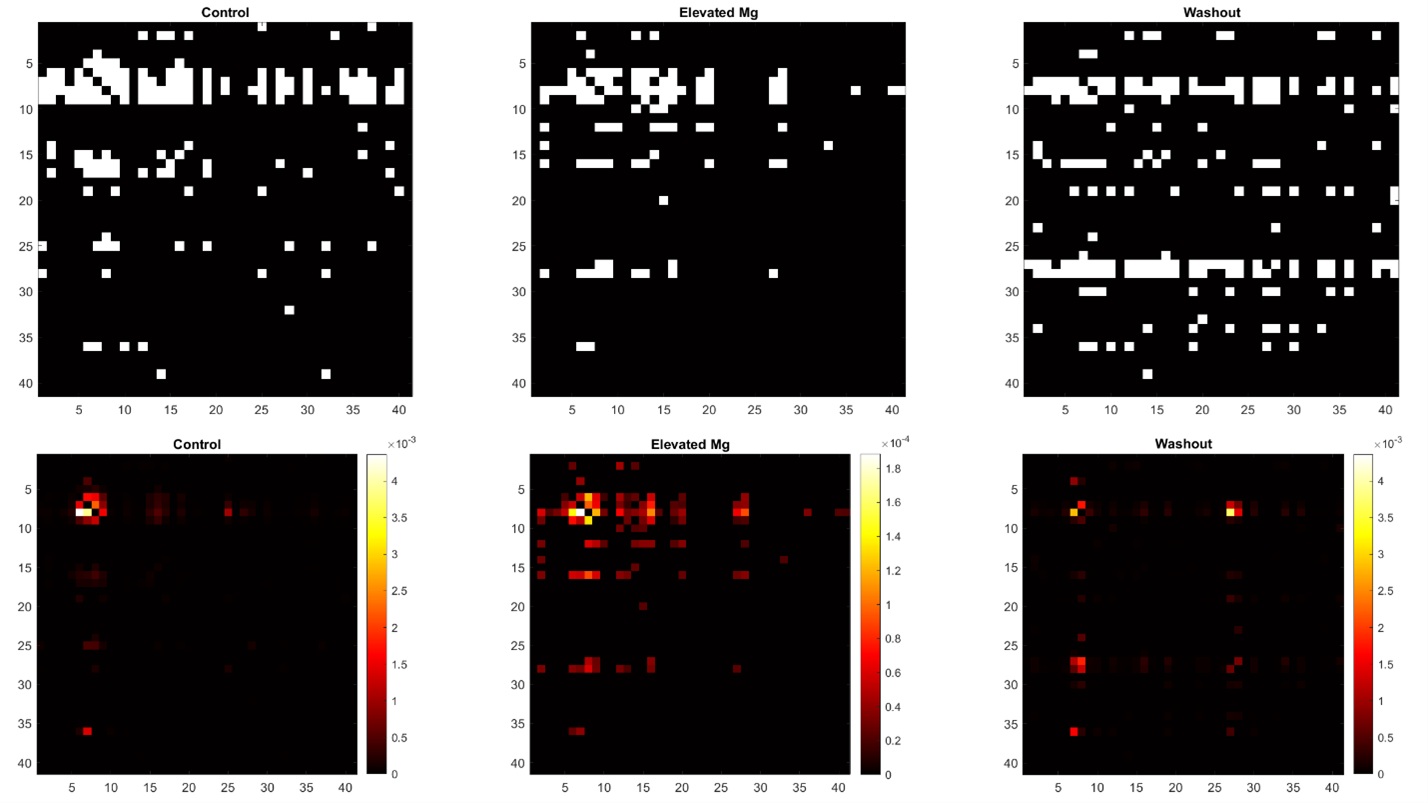

### Supplemental Figure 2

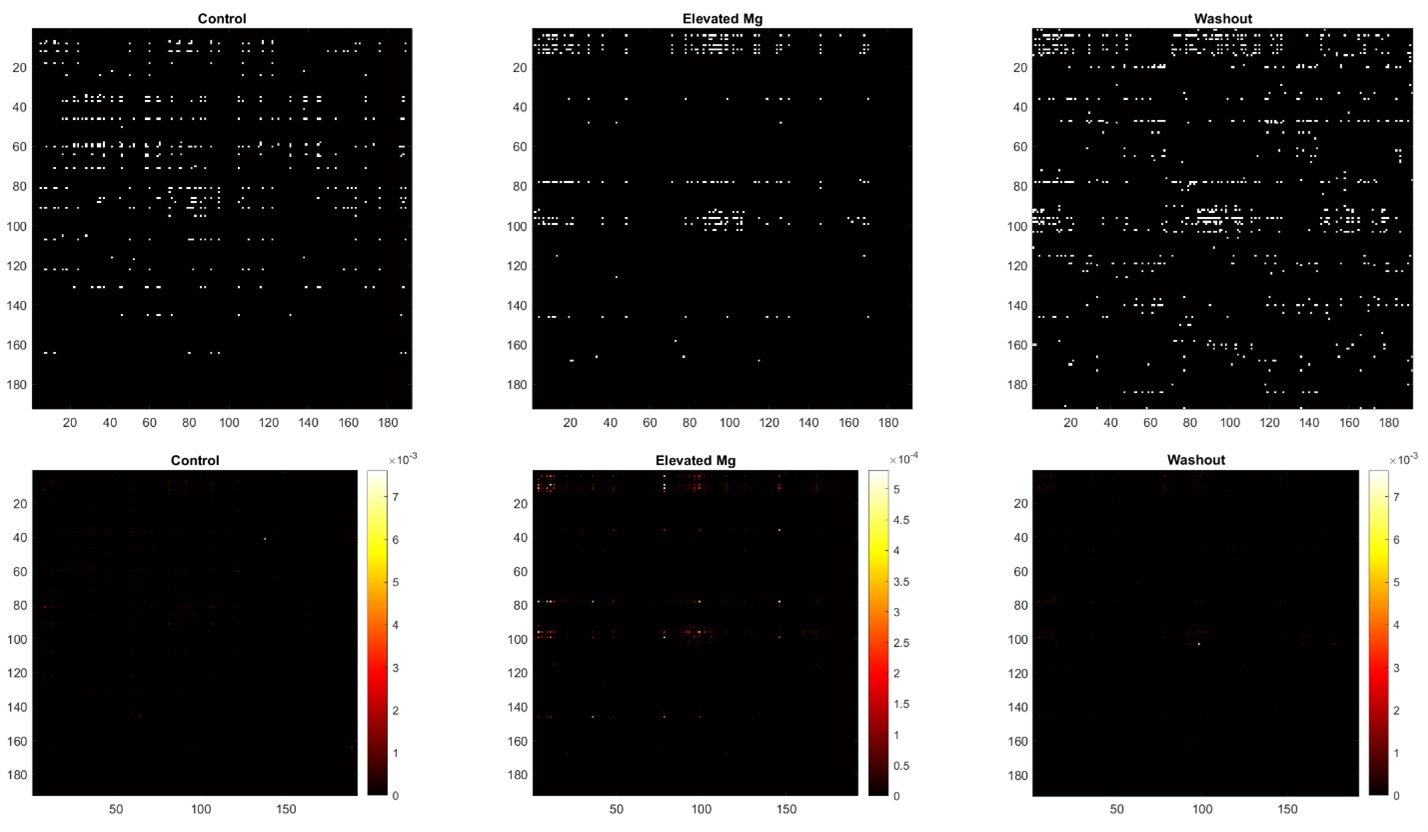
